# Assembly-free rapid differential gene expression analysis in non-model organisms using DNA-protein alignment

**DOI:** 10.1101/2021.04.23.441097

**Authors:** Anish M.S. Shrestha, Joyce Emlyn B. Guiao, Kyle Christian R. Santiago

## Abstract

**Background:** RNA-seq is being increasingly adopted for gene expression studies in a panoply of non-model organisms, with applications spanning the fields of agriculture, aquaculture, ecology, and environment. For organisms that lack a well-annotated reference genome or transcriptome, a conventional RNA-seq data analysis workflow requires constructing a de-novo transcriptome assembly and annotating it against a high-confidence protein database. The assembly serves as a reference for read mapping, and the annotation is necessary for functional analysis of genes found to be differentially expressed. However, assembly is computationally expensive. It is also prone to errors that impact expression analysis, especially since sequencing depth is typically much lower for expression studies than for transcript discovery.

**Results:** We propose a shortcut, in which we obtain counts for differential expression analysis by directly aligning RNA-seq reads to the high-confidence proteome that would have been otherwise used for annotation. By avoiding assembly, we drastically cut down computational costs – the running time on a typical dataset improves from the order of tens of hours to under half an hour, and the memory requirement is reduced from the order of tens of Gbytes to tens of Mbytes. We show through experiments on simulated and real data that our pipeline not only reduces computational costs, but has higher sensitivity and precision than a typical assembly-based pipeline. A Snakemake implementation of our workflow is available at: https://bitbucket.org/project_samar/samar

**Conclusions:** The flip side of RNA-seq becoming accessible to even modestly resourced labs has been that the time, labor, and infrastructure cost of bioinformatics analysis has become a bottleneck. Assembly is one such resource-hungry process, and we show here that it can be avoided for quick and easy, yet more sensitive and precise, differential gene expression analysis in non-model organisms.

## Background

RNA-seq has become the principal technique for measuring variation of genome-wide gene expression levels across conditions [1]. Differential expression analysis usually begins by mapping RNA-seq reads to either a reference genome or transcriptome sequence. On one hand, accurate genome annotation has not kept up with the increase in sequence data [2]. Consequently, well-annotated and high-quality reference sequences are available for only a handful of model organisms. On the other hand, driven by declining costs, RNA-seq is becoming increasingly accessible to labs with modest resources; and as a result, it is being employed on an ever-expanding catalog of non-model organisms, pervading the fields of agriculture, aquaculture, ecology, and environment. A very short list of recent studies include: environmental stress response in sea-trout [3], coral [4], ryegrass [5], pigeonpea [6], tiger barb [7]; immune response to parasites and pathogens in guppy [8], eel [9], silkworm [10], peanut [11], sunflower [12]; mechanisms of phenotypic divergence in hares [13], bats [14], grass carps [15]; effect of diet in the growth and development in shrimp [16], yellow perch [17], mandarin fish [18], grenadier anchovy [19], catfish [20], tilapia [21], bass [22]. It is only likely that RNA-seq will continue to rapidly proliferate while high-quality reference databases grow at a slow pace.

The conventional strategy to adapt standard reference-based RNA-seq analysis workflows to the case of nonmodel organisms, has been to first compute a de-novo transcriptome assembly by pooling all reads and to annotate the assembly against a high-confidence protein database. Since assemblers typically do not provide a read-to-contig mapping, the subsequent step is to map the reads to the assembly. This is followed by a quantification step in which reads mapping to each contig are counted. The count data is used for statistical testing of differential expression. Since the number of differentially expressed genes tends to be quite large, inference of biological function is done computationally by using the annotations to perform GO-term enrichment or pathway analysis.

A major drawback of de-novo assembly is that it requires massive computational resources. In most cases, the goal is to characterize the expression profile of the protein-coding fraction of the transcriptome, and not necessarily to obtain an assembly. Accordingly, samples are sequenced at a much lower depth than would be required for a reliable assembly [23]. Assembly errors such as over-extension, fragmentation, and incompleteness of contigs can adversely impact downstream expression analysis [24, 25]. Furthermore, assemblers tend to over-estimate the number of isoforms/contigs per gene, which introduces complications for statistical test of differential expression as well as interpretation of results since many genes could appear multiple times in the final result. These issues have motivated supplementary measures such as clustering contigs [26] or aggregating expected read counts of contigs which map to the same reference gene [27], prior to statistical testing of differential expression. Finally, in special cases such as comparison of gene expression across species, it might not even be reasonable to compute a single assembly.

We provide an alternative strategy that circumvents the need for assembly and annotation. The first step of our proposed pipeline uses LAST [28, 29] to directly align RNA-seq reads to the high-confidence protein set which would otherwise have been used for annotation. This is followed by a simple counting step that employs a traditional rescue strategy to resolve multimaps [30]. The counts can be fed into standard count-based differential gene expression analysis tools, e.g. DESeq2 [31]. Our main proposition here is that since functional analyses in non-model organisms rely on a database of homologous proteins in order to draw conclusions, it might be more reasonable to directly allocate reads to those homologous proteins using DNA-protein alignment, instead of introducing an error-prone yet computationally heavy intermediary step of assembly.

The main and obvious advantage of our method is that it drastically brings down computational costs. For example, for a typical RNA-seq study containing 2 groups of 3 replicates each and 20 million paired-end reads per replicate, our approach takes under half an hour, whereas computing an assembly would take several tens of hours. Additionally we show, through experiments on simulated and real RNA-seq datasets, that our method is more accurate in identifying differentially expressed genes than an assemblymapping-quantification pipeline. Another advantage is that it is easier to interpret results, as each homologous gene is reported as differentially expressed or not, along with associated statistical measures. In contrast, with assembly-based pipelines, there might be a need to consolidate results across several fragmented contigs. Furthermore, reference proteomes, for example in UniProt, come with GO annotations, allowing for a straightforward transition to downstream functional analysis; whereas with assembly-based pipelines, there is a need to post-process the multiple local alignments that might be reported by the annotation software for each contig.

## Implementation

### Implementation of our proposed method

We have implemented our proposed strategy as a Snakemake pipeline [32], which is available at https://bitbucket.org/project_samar/samar.

The first step in the workflow aligns RNA-seq reads to a reference set of proteins. For this we use the DNA-protein alignment feature of LAST [28, 29, 33]. We chose LAST over numerous other aligners capable of DNA-protein alignment – BLASTX [34] being a prominent example – for its unique combination of features. It scales well to high-throughput sequencing data. The probabilistic framework for incorporating paired information from paired-end reads, which was originally designed for read-to-genome alignment [35], works out of the box for the case of read-to-proteome alignment. It allows training the substitution matrix and gap penalties to reflect the sequence divergence between the (translated) RNA-seq reads and the reference proteome [36].

In the second step, from the alignments produced by LAST, we compute counts of reads originating from each entry in the reference. This is not trivial due to multi-mapping, an issue that becomes more pronounced when the reference contains isoforms with high sequence similarity. We employ the simple strategy of rescuing multi-mapping reads proposed in [30], where the contribution of a read mapping to several protein sequences is distributed based on their read coverage estimated from uniquely mapping reads. Suppose the reference is a set of protein sequences indexed by *P* = {1, 2, …, *n*}. The counting proceeds in two passes. In the first pass, we obtain the count of reads aligning uniquely to sequence *i*, normalized by the length of *i* covered by the uniquely mapping reads. Let us denote this normalized count by *u*_*i*_. In the second pass, for each read multi-mapping to a subset *P′* ⊆ *P*, we update the count of sequence *i* ∈ *P′* in proportion to *u*_*i*_, i.e. to the current count of sequence *i*, we add *c*_*i*_, where

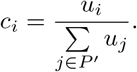

If the denominator is zero, we distribute the count evenly among *P′*.

The counts obtained in the second step can be fed directly to count-based differential gene expression analysis tools. In our pipeline, we use DESeq2 [31].

### Benchmarking

We perform benchmarking on both real and simulated RNA-seq read datasets from the protein-coding transcriptome of the fruit fly *D. melanogaster*. We chose this extensively studied transcriptome since the transcripts and protein products associated with each gene is known for a huge number of protein-coding *D. melanogaster* genes. Such a mapping between Flybase ID and UniProt ID can be obtained from Ensembl, UniProt, Flybase [37], etc., and can be used for evaluation of our workflow.

The simulated RNA-seq read dataset was generated as follows. We downloaded the transcripts of proteincoding genes from the fruit fly assembly BDGP6.28 obtained from Ensembl Genes 101. After removing sequence duplicates and transcripts with no corresponding protein entries, there were 28,692 transcripts of 13,320 genes. From this transcriptome, we simulated 2 groups of RNA-seq reads with 3 replicates per group using Polyester [37]. In the first group, the mean expression levels of the transcripts were set to be proportional to the FPKM values computed from an arbitrarily chosen poly-A+ enriched real RNA-seq data (ArrayExpress E-MTAB-6584). The FPKM values were estimated using RSEM [38] on Bowtie2 [39] alignments of the reads to the transcriptome. In the second group, a subset of roughly 30% of the transcripts were set to be differentially expressed at varying levels of up- and down-regulation: 1.5, 2, and 4-fold. The transcripts were chosen by randomly selecting genes and setting only the highest expressing isoform to be differentially expressed. Since inference of differential expression is typically done at the gene level, having at most one isoform to differentially expressed simplifies the evaluation process [40] as we can define a gene to be differentially expressed if one of its transcripts was differentially expressed. In fact, it might not be too far from reality as it has been shown that most highly expressed protein-coding genes have a single dominant isoform [41]. Each read set had roughly 20 million pairs of 100 bp reads with mean fragment length of 250 bp.

## Results

### DNA-protein alignment attains similar performance to using a transcriptome reference

We first demonstrate, under ideal conditions, the soundness of our idea of aligning RNA-seq reads to a proteome reference for differential gene expression analysis, by comparing our performance to that of the traditional case of using an established transcriptome reference.

We aligned the fruit fly simulated RNA-seq reads (described earlier in the Implementation section) to the UniProt *D. melanogaster* proteome UP0000 00803, which contains 1 representative protein sequence per gene, and fed the counts obtained by our method to DESeq2 [31] for differential analysis. To serve as baseline for comparison, we additionally ran a typical pipeline consisting of Bowtie2 [39] for read alignment to the *D. melanogaster* transcriptome, followed by RSEM for transcript quantification, tximport [40] for gene-level aggregation of counts, and finally DESeq2 [31] for differential analysis at the gene level. Details of the two pipelines are provided in the Supplementary Material.

We evaluated the two approaches based on their recall and precision in predicting differentially expressed (DE) genes. Recall is the proportion of actual DE genes that were correctly predicted to be DE, and precision is the proportion of predicted DE genes that were actually DE. We require that the direction of fold-change (up/down) match between the ground truth and prediction to be classified as a correct prediction. To compute recall and precision of our method, we mapped our predictions from the set of proteins to the set of genes using the mapping between UniProt protein ID and FlyBase gene ID obtained from Ensembl.

Figure 1(left) shows the Precision-Recall curves obtained by varying the false discovery rate (FDR) threshold of DESeq2, and Figure 1(right) shows the distribution of the estimated log fold-change at an FDR threshold of 0.1. There is almost no difference between the two approaches in the ability to detect DE genes and in the trend of under- or over-estimation of true fold change. This result demonstrates that for differential gene expression analysis, there is no performance degradation when aligning RNA-seq reads to a proteome reference, even though we are effectively only using reads from the coding region of transcripts.

**Figure 1:**
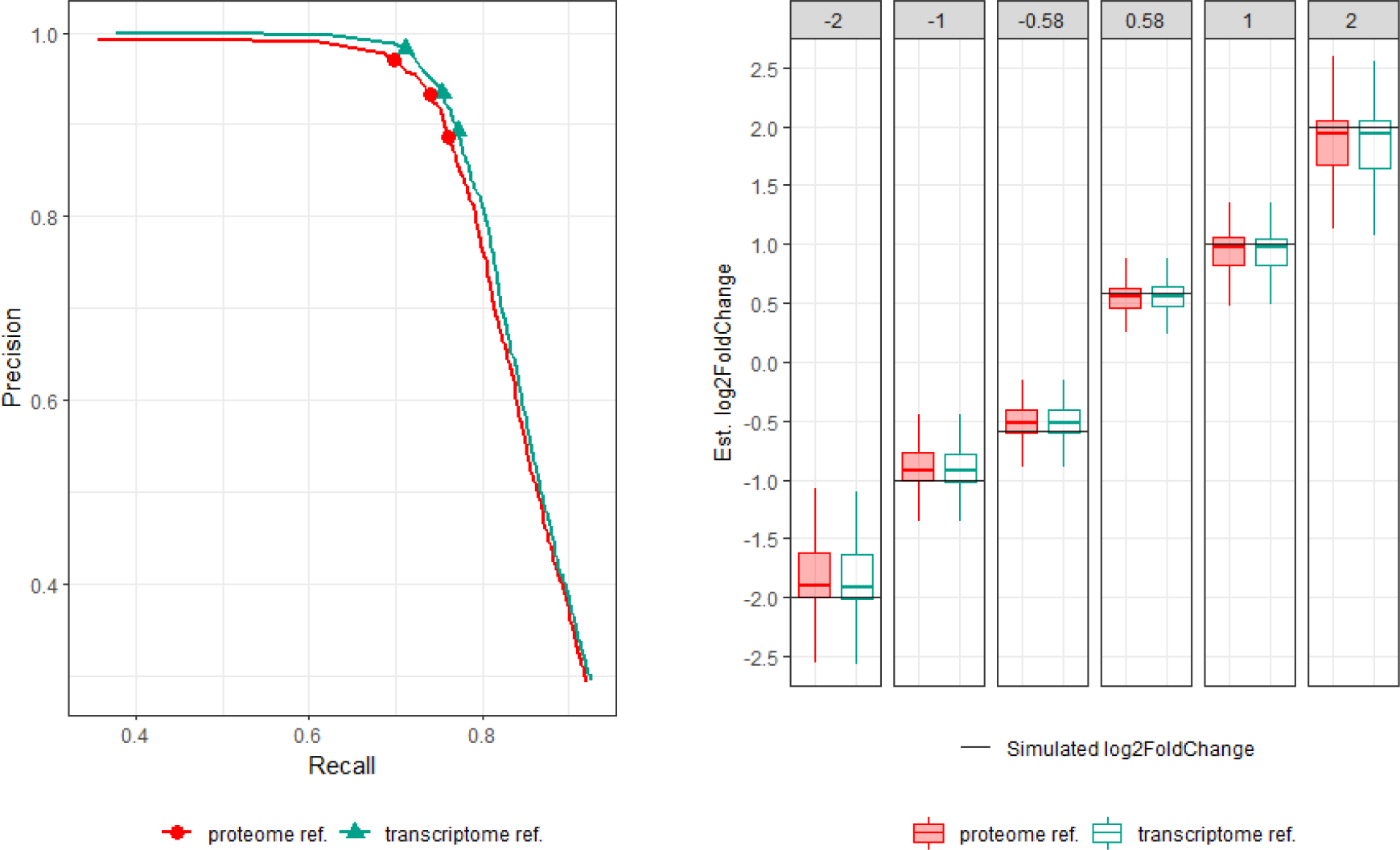
(Left) Precision-recall curves of our method using the *D. melanogaster* proteome reference and the Bowtie2-RSEM-DESeq2 pipeline using its transcriptome reference. The three open dots in each curve correspond to setting the FDR threshold of DESeq2 to 0.01, 0.05, and 0.1. (Right) Log fold change of true positive DE genes estimated by DESeq2 at FDR threshold of 0.1, compared against the 6 simulated log-fold change levels.

### In the absence of a close reference, our method outperforms a typical assembly-based approach

Next we simulated the scenario faced in the case of non-model organisms by pretending that the *D. melanogaster* reference sequences are not available, and that the closest species with a well-annotated reference proteome is a distant relative, the mosquito *Anopheles gambiae*. The two species are of the same order Diptera with their lineages thought to have separated roughly 250 million years ago [42]. The evaluation process described below uses the mosquito proteome as reference; but to calibrate the effect of the degree of evolutionary and sequence divergence, we repeated the process described below with reference proteomes of closer relatives of *D. melanogaster*: *D. ananassae* and *D. grimshawi*.

We compared our performance to that of a typical assembly-based pipeline consisting of: Trinity [43] for de-novo transcriptome assembly, followed by Bowtie2 for mapping the reads to the assembly, RSEM for counting, tximport for gene-level aggregation using the gene-to-transcript mapping provided by Trinity, and finally DESeq2 for differential analysis. We used the Dammit pipeline [44] to annotate the assembly against the mosquito proteome. Details of the two pipelines are provided in the Supplementary Material.

To facilitate the comparison, we obtained a precomputed orthology map between *A. gambiae* and *D. melanogaster*, from the website of InParanoid [45]. Consider a *D. melanogaster* protein-coding gene *g*, and let *F*_*g*_ be the set of protein products of *g*. For a *D. melanogaster* protein *f*, let *O*_*f*_ be the set of mosquito proteins in the same ortholog group as *f*. We associate with *g* the set *M*_*g*_ of mosquito proteins *m* such that *m* ∈ *O*_*f*_ for some *f* ∈ *F*_*g*_.

We computed recall and precision of our method as follows. An actual up-regulated (down-regulated) *D. melanogaster* DE gene *g* was defined as correctly predicted if there was at least one protein in *M*_*g*_ that was predicted to be up-regulated (down-regulated). Recall was defined as the number of correctly predicted DE genes divided by the number of actual DE genes. Precision was defined as the number of correctly predicted DE genes divided by the number of genes *g* for which at least one protein in *M*_*g*_ was predicted to be DE.

We computed recall and precision of the assembly-based approach as follows. For a Trinity gene *t*, let *D*_*t*_ be the set of mosquito proteins that Dammit assigned to the isoforms of *t* (if there were multiple alignments for an isoform, we kept only one with the lowest Evalue). With an actual *D. melanogaster* gene *g*, we associated a set *T*_*g*_ of Trinity genes, where *t* ∈ *T*_*g*_ if *D*_*t*_ ∩ *M*_*g*_ = ∅. An actual up-regulated (down-regulated) DE *D. melanogaster* gene *g* was defined to be correctly predicted if there was at least one up-regulated (downregulated) Trinity gene in *T*_*g*_. Recall was defined as the number of correctly predicted DE genes divided by the number of actual DE genes. Precision was defined as the number of correctly predicted genes divided by the number of genes *g* for which at least one gene in *T*_*g*_ was predicted to be DE.

The definitions of recall and precision are necessarily slightly different for the two approaches. Our hope is that they convey a similar meaning – that an actual *D. melanogaster* DE gene is represented by a set of orthologous mosquito proteins (in our method) or by a set of Trinity genes for which there was an annotation to an orthologous mosquito protein (in the assembly-based method), and that the gene is considered to be correctly predicted if at least one of the representatives are predicted to be DE.

Figure 2 shows the precision and recall of our method and the assembly-based approach when using the mosquito reference proteome. It also contains the PR-curves when using the *D. ananassae* and *D. grimshawi* reference proteomes. The curves were obtained by varying the FDR threshold of DESeq2. When using the two *Drosophila* reference proteomes, the performance of our method varied slightly, but in both cases, outperformed the assembly-based approach. When using the *A. agambiae* reference, recall was lower for both methods, mainly because the orthology map contains only 60% of the fruit fly proteins – there were 7341 ortholog clusters involving 7863 fruit fly proteins and 8090 mosquito proteins.

**Figure 2:**
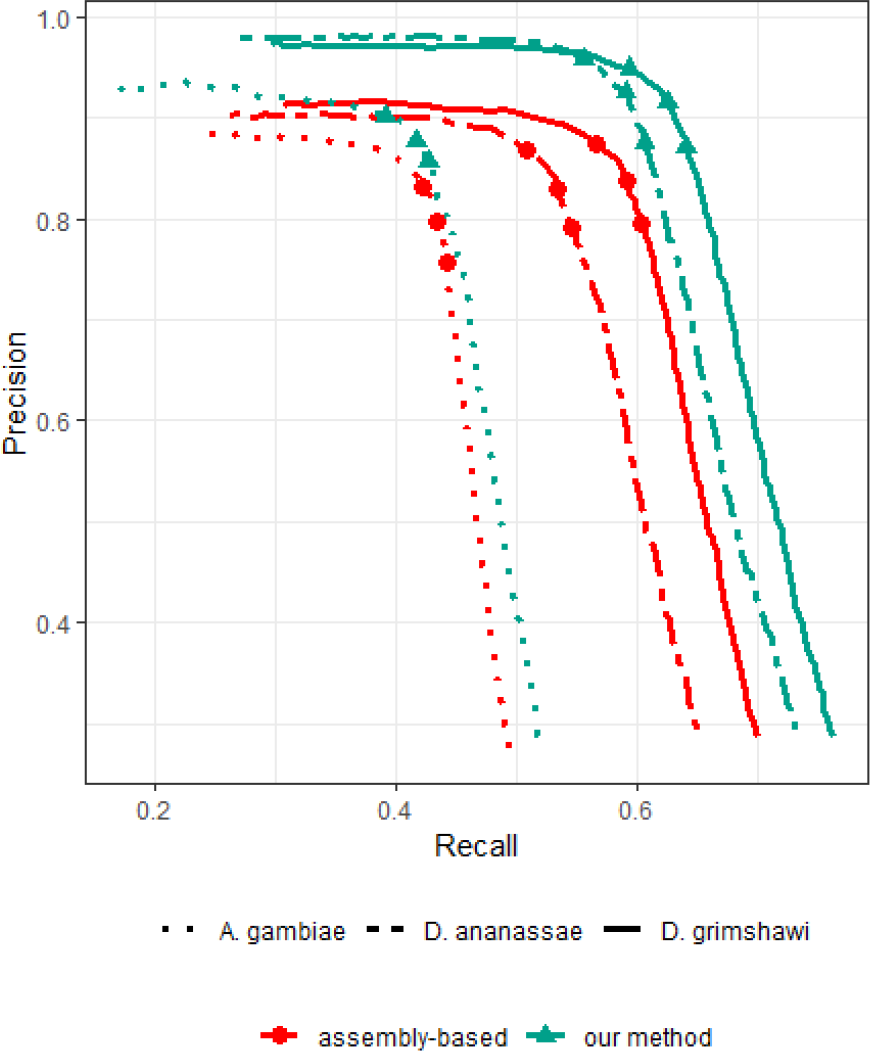
Precision-Recall curves of our method and the assembly-based pipeline, when using reference proteomes of close relatives (*D. ananassae* and *D. grimshawi*) and a distant relative (*A. gambiae*). The three open dots in each curve correspond to setting the FDR values of 0.01, 0.05, and 0.1.

Overall, across any setting of FDR threshold or any choice of a reference proteome, our approach outperformed the assembly-based approach.

So far, to compute recall and precision of the assembly-based approach, we used all the alignments reported by the Dammit pipeline, even including many short local alignments. It is not uncommon in practice to filter short alignments. We repeated the analysis by keeping only those alignments predicted by the Dammit pipeline that covered at least 50% of a contig. The precision-recall curves for this cases is shown in Figure 3, which shows a significant drop in recall of the assembly-based approach.

**Figure 3:**
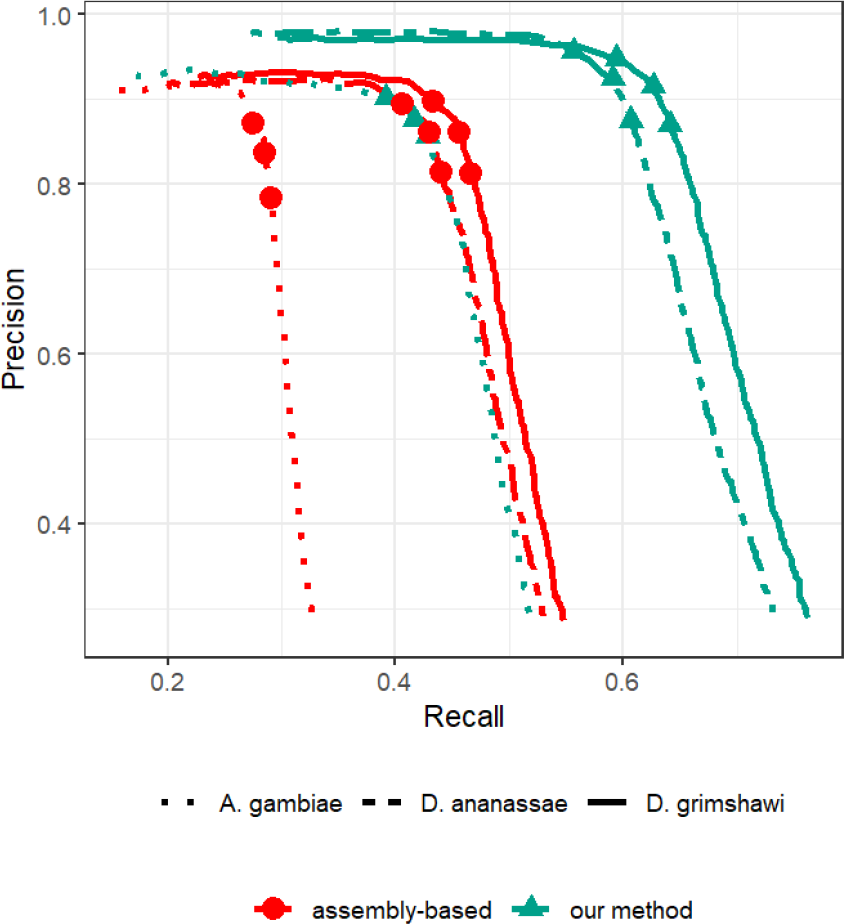
Precision-Recall curves of our method and the assembly-based pipeline, when using reference proteomes of close relatives (*D. ananassae* and *D. grimshawi*) and a distant relative (*A. gambiae*), and with the alignments produced by Dammit which covered less than 50% of the length of the contig removed. The three open dots in each curve correspond to setting the FDR values of 0.01, 0.05, and 0.1.

### With real data too, our method outperforms the assembly-based approach

We applied our pipeline and the assembly-based approach to a recently published real RNA-seq dataset ArrayExpress E-MTAB-8090 ERR3393437–42. The dataset contains RNA-seq reads of the hemocyte tissue of *D. melanogaster* samples with and without injury, with 3 replicates for each condition. After cleaning and trimming low-quality reads using fastp [46], there were roughly 110 million pairs of reads.

Continuing with the assumption that no reference sequences are available for *D. melanogaster*, we applied our pipeline and the assembly-based pipeline as in the previous section using the mosquito and two *Drosophila* reference proteomes. Since we do not know the ground truth for this dataset, to serve as baseline, we additionally ran the Bowtie2-RSEM-DESeq2 pipeline using *D. melanogaster* reference transcriptome. To be able to compare the DE call sets, we mapped our predicted DE genes to the *D. melanogaster* gene names using the same Inparanoid orthology maps as before.

Intersection of the sets of DE genes obtained from the two approaches and the baseline are shown in Figure 4. At FDR threshold of 0.01, there were 104 genes identified as DE by the baseline method. Based on the observation from Figure 1 that the Bowtie2-RSEM-DESeq2 pipeline has high precision at FDR 0.01, let us assume that all of these baseline calls are correct and that they constitute the empirical ground truth. At the same FDR threshold, when using the *D. ananassae* reference proteome, our method was slightly more sensitive than the assembly-based approach, being able to predict 68 out of the 104 baseline DE genes, compared to 58 by the assembly-based approach. Our method was also slightly more precise, with the 68 calls corresponding to roughly 78% of the calls, compared to 74% for the assembly-based method. This is in line with observation from Figure 2 that our method has slightly better sensitivity and precision than the assembly-based approach. Similar results were obtained when using the *D. grimshawi* reference proteome.

**Figure 4:**
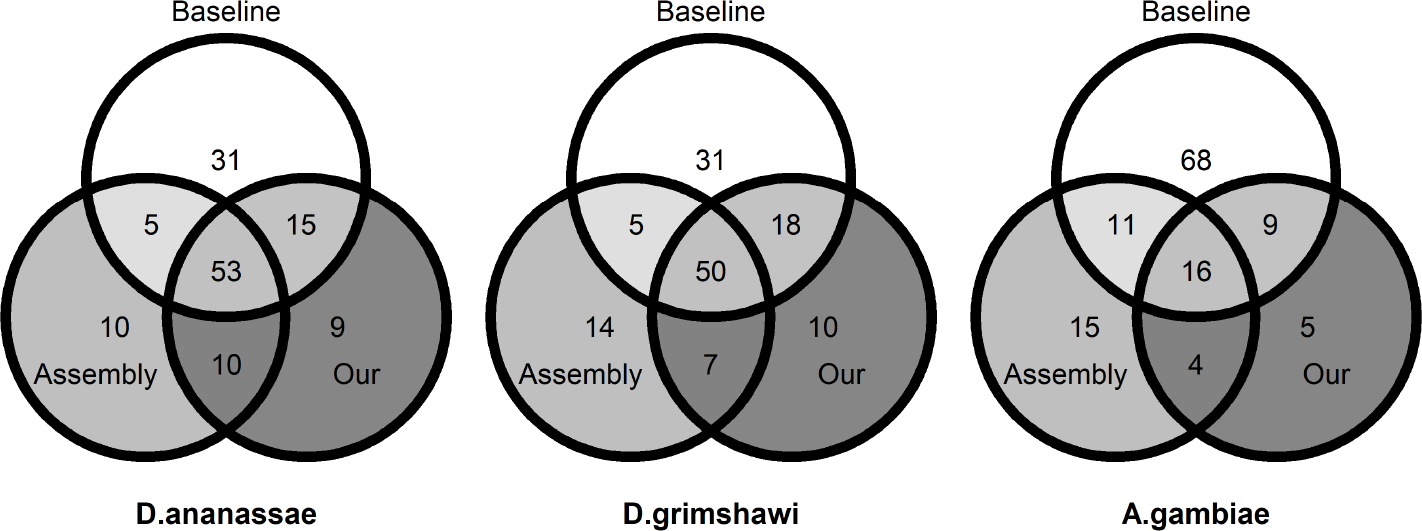
For the three reference proteomes, Venn diagrams showing the intersections among the **Baseline** set consisting of DE genes called by the baseline approach of Bowtie2-RSEM-DESeq2 using the *D. melanogaster* reference transcriptome, **Our** set consisting of *D. melanogaster* genes to which the DE genes called by our approach mapped to, and (3) **Assembly-based** set consisting of *D. melanogaster* genes to which Trinity DE genes mapped to. FDR threshold of 0.01 was used for all three approaches.

When using the *A. gambiae* proteome, there is a significant decrease in the size of the overlaps between the baseline and the two approaches, consistent with the drop in sensitivity observed in Figure 2. The two approaches are similarly sensitive (25 calls by our approach vs. 27 by assembly-based) while our method is more precise (25 out of 34 calls by our approach vs. 27 out of 46 calls by assembly-based).

### Avoiding assembly dramatically reduces running time and memory usage

All the experiments above were carried out on a system with Intel Xeon Silver 4114 Processor with 10 cores and 20 threads. For the real dataset E-MTAB-8090 which contains roughly 110 million pairs of cleaned reads, de-novo assembly alone took more than 24 hours. In contrast, DNA-protein alignment, which is the most compute-intensive part of our pipeline, takes less than 20 minutes per sample containing roughly 20 million pairs of reads, using 20 threads. While the denovo assembly had a massive peak memory usage of ∼ 65 Gbytes, the memory requirement of our method is dominated by the size of the proteome index, which was just ∼ 33 Mbytes for the mosquito proteome. In general, the index size is roughly 5 × *n* bytes, where *n* is the length of the proteome.

## Discussion

### Summary of results

We have shown that aligning RNA-seq reads to a proteome reference followed by a simple counting procedure provides an extremely fast and light-weight alternative to the current resource-intensive assemblyand-annotation based approach for differential gene expression analysis. We have shown through experiments on simulated and real datasets that our approach is more sensitive and precise than the assembly-based approach.

### Isoform-level quantification

In this paper, we focused on differential expression analysis at the gene level, as it has been shown that it is advantageous to perform statistical inference of differential expression at the gene level even if the quantification is done at the transcript level [40]. Since we used reference sets with only 1 protein entry per gene, our counts were automatically at the gene level. However, it is also possible to get isoform-level counts and aggregate the counts at the gene level for differential analysis. We saw no loss of performance with this approach (Supplementary Material). An advantage of isoform-level counts is that it can be used for other kinds of statistical tests such as differential usage of isoforms across conditions. This is akin to the differential transcript expression/usage studies.

### Choice of reference

Our results suggest – not surprisingly – that the choice of reference has a huge influence on the outcome of differential expression analysis, since a distant reference means fewer reads are aligned (correctly). One source to find a closest possible proteome is the UniProt Reference Proteome database. This database currently contains almost 20,000 proteomes of organisms which are relatively well-studied and “provide broad coverage of the tree of life” [47].

Apart from single-species reference proteomes, it is also common to use cross-species proteins sets such as Swiss-Prot for annotating transcriptome assemblies. In theory, our method can also use Swiss-Prot as reference. However, Swiss-Prot is extremely redundant due to presence of orthologous proteins, which can needlessly aggravate the issue of multi-mapping. To use Swiss-Prot as reference, it is advisable to remove sequence redundancies by using tools such as CDHIT [48] or MMSeq2 [49] and by selecting a subset of Swiss-Prot based on taxa.

### Room for improvement

Currently we use a simple technique of rescuing multimapping reads. It would be interesting to explore a more sophisticated way to handle multi-mapping issues similar to a statistical model in RSEM.

LAST currently does not handle quality data present in the fastq records during alignment, and as far as we know, nor do other DNA-protein aligners. It is an interesting open problem to investigate if incorporating the quality data improves alignment accuracy, not just in this application to RNA-seq data analysis but to other applications of DNA-protein alignment.

### Long reads

This paper is focused on short-read datasets, since from our cursory literature search in the Introduction section, it appears that long-read technologies are currently not widespread as in the applied fields that deal with non-model organisms. Theoretically, the core idea of DNA-protein alignment carries over just as well to long reads. Longer sequences can potentially improve accuracy as it would be easier to disambiguate counts among paralogous genes. However, application to long reads warrants a separate benchmarking process as one needs to account for error profiles and error rates characteristic to long-read technologies.

## Conclusions

With RNA-seq becoming accessible to even labs with limited resources, the time, labor, and infrastructure cost of bioinformatics analysis has grown. Transcriptome assembly is one such resource-hungry process, which takes several tens of hours on typical datasets, even on high-performance computing systems. For many labs, such requirements can impose a serious bottleneck. By avoiding assembly, our pipeline allows for quick and easy, yet more sensitive and precise, analysis of differential gene expression in non-model organisms.

## Availability and requirements

Project Name: SAMAR (Speedy, Assembly-free Method to Analyze RNA-seq expression data)

Project Home page: https://bitbucket.org/project_samar/samar

Operating systems: Linux or MacOS

Programming language: Snakemake workflow management system

Other requirements: Conda. All dependencies automatically handled by Snakemake.

License: MIT License

## Abbreviations

GO: Gene Ontology FDR: False discovery rate
DE: Differentially expressed

## Declarations

### Ethics approval and consent to participate

Not applicable.

### Consent for publication

Not applicable.

### Availability of data and materials

Our software is available at

https://bitbucket.org/project_samar/samar.

Scripts used in this paper for generation of simulated data and benchmarking study is available at: https://bitbucket.org/project_samar/benchmarking.

Datasets for benchmarking were obtained from public repositories: transcripts of protein-coding genes were obtained from the fruit fly assembly BDGP6.28 in Ensembl Genes 101; reference proteomes of *D. melanogaster* (UP000000803), *D. grimshawi* (UP000001070), *D. ananassae* (UP000007801), and *A. gambiae* (UP000007062) were obtained from UniProt.

### Authors’ contributions

AMSS planned the study. AMSS and JEBG performed the benchmarking study. AMSS, JEBG, and KCRS wrote the software. AMSS wrote the manuscript with inputs from all authors. All authors read and approved the final manuscript.

## Acknowledgements

We are grateful to Martin Frith for his advice on DNA-protein alignment and LAST usage, and for providing helpful comments on the manuscript. We thank Hugues Richard for suggesting many improvements to the manuscript.

## Additional Files

Additional file 1 — Supplementary Material

